# Mitochondrial Ca^2+^ uniporter (MCU) variants form plasma-membrane channels

**DOI:** 10.1101/2023.07.31.551242

**Authors:** Iuliia Polina, Jyotsna Mishra, Michael W Cypress, Maria Landherr, Nedyalka Valkov, Isabel Chaput, Bridget Nieto, Ulrike Mende, Peng Zhang, Bong Sook Jhun, Jin O-Uchi

## Abstract

MCU is widely recognized as a responsible gene for encoding a pore-forming subunit of highly mitochondrial-specific and Ca^2+^-selective channel, mitochondrial Ca^2+^ uniporter complex (mtCUC). Here, we report a novel short variant derived from the MCU gene (termed MCU-S) which lacks mitochondria-targeted sequence and forms a Ca^2+^- permeable channel outside of mitochondria. MCU-S was ubiquitously expressed in all cell-types/tissues, with particularly high expression in human platelets. MCU-S formed Ca^2+^ channels at the plasma membrane, which exhibited similar channel properties to those observed in mtCUC. MCU-S channels at the plasma membrane served as an additional Ca^2+^ influx pathway for platelet activation. Our finding is completely distinct from the originally reported MCU gene function and provides novel insights into the molecular basis of MCU variant-dependent cellular Ca^2+^ handling.

## Introduction

The discovery of the molecular identity of the pore-forming subunit of the mitochondrial Ca^2+^ (mtCa^2+^) uniporter complex (mtCUC) termed MCU (originally known as CCDC109A) in 2011 (Baughman et al, 2011, De Stefani et al, 2011) has been providing new possibilities for applying genetic approaches to precisely understand the mtCa^2+^ handing mechanism and its impact to the mitochondrial and cellular functions. Indeed, rapid progress was attained in characterizing the role of MCU via genetic knockout/overexpression studies of MCU both in both cell systems and *in vivo* (Alevriadou et al, 2021, Jhun et al, 2016, Mammucari et al, 2018). Although the alteration of mtCa^2+^ via MCU is frequently observed in human diseases with disrupted energy metabolism (Calvo-Rodriguez et al, 2020, Ghosh et al, 2020, Zaglia et al, 2017), it is still not clear how MCU protein expression and mtCUC function are regulated by the MCU gene under both physiological and pathological conditions. MCU was originally considered to be a highly mitochondria-specific and Ca^2+^-selective channel in the IMM with other subunits including MICU families, EMRE and MCUb (Alevriadou et al, 2021, Jhun et al, 2016, Mammucari et al, 2018), mainly because the original/conventional MCU (“long form” MCU, here renamed MCU-L) possesses a predicted cleavable N-terminal mitochondrial targeted sequence (MTS) (Baughman et al, 2011, De Stefani et al, 2011). Data sets published prior to 2011 had predicted the existence of additional, still uncharacterized human and mouse MCU cDNAs (Carninci et al, 2005, Kawai et al, 2001, Okazaki et al, 2002, Strausberg et al, 2002), but the expression patterns and functional roles of these potential MCU variants have not yet been reported. Here, we report that the CCDC109A gene encodes ion channels that are located both inside and outside the mitochondria, especially in human platelets to form Ca^2+^-permeable channel at the plasma membrane (PM), which greatly expands and revises the established concept of MCU (Baughman et al, 2011, De Stefani et al, 2011) as a mitochondria-specific Ca^2+^ channel.

## Results

### Detection of short MCU transcript variants

MCU (originally known as CCDC109A) genes in humans and mice consist of 10 and 9 exons, respectively. Based on a thorough search of online databases and published data sets of complete sequences for full-length, uncharacterized human and mouse cDNAs (Carninci et al, 2005, Kawai et al, 2001, Okazaki et al, 2002, Strausberg et al, 2002), we identified 3 and 2 potential protein-coding transcripts from CCDC109A gene, respectively (**Fig. 1**). MCU-L in both mice and humans is originally reported as the transcript encoding the pore for mtCUC (Baughman et al, 2011, De Stefani et al, 2011).

**Fig. 1.**
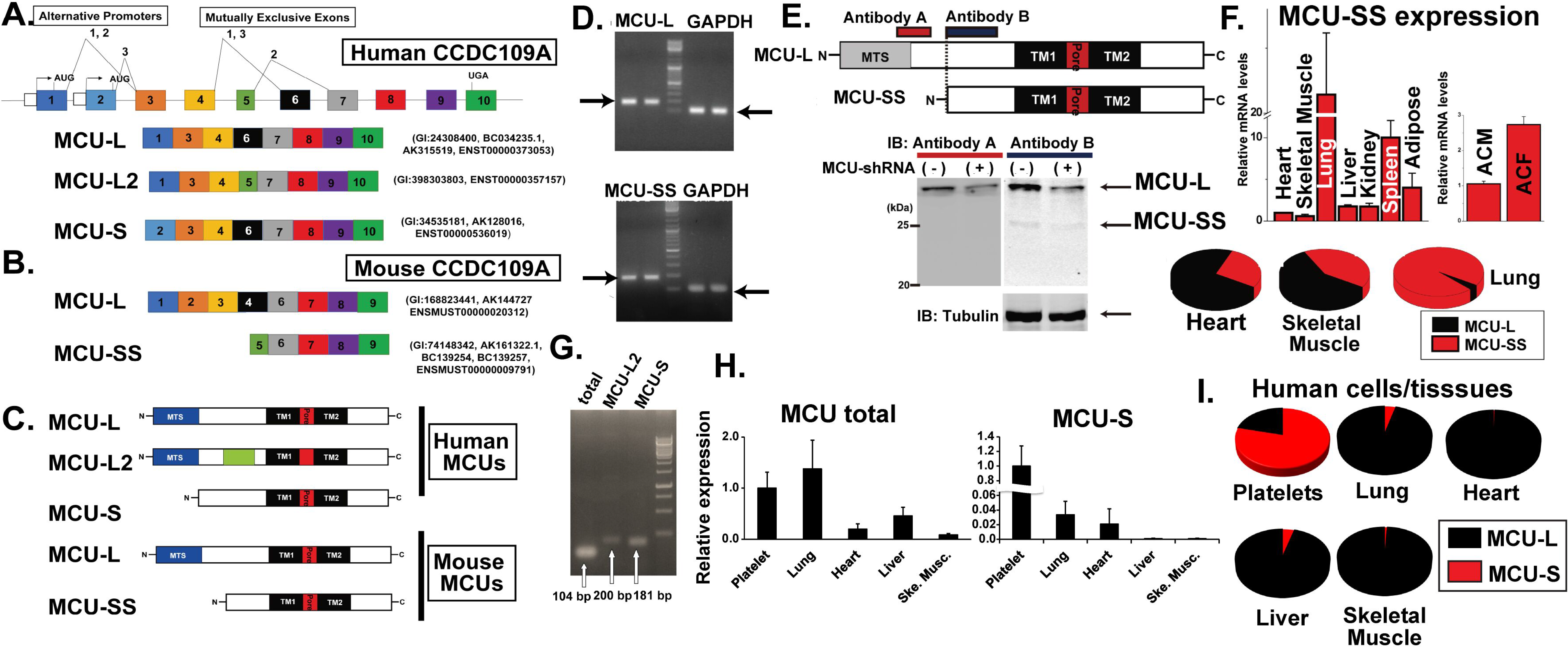
Detection of MCU variants in human and mouse cells/tissues. **A.** Structural organization of predicted human MCU (CCDC109A) variants. The colored cassettes stand for the exons and the blank cassettes show the predicted promoters. **B.** Predicted mouse MCU (CCDC109A) variants. **C.** Schematic protein structures of human and mouse MCU variants (See also Supplementary Figure 1). TMD, transmembrane domain. **D.** PCR of mouse MCU variants. cDNA was created from mRNA extracted from C2C12 cells. The 1.8 kb fragments were amplified by a pair of MCU variant-specific primers. GAPDH was used as a control. **E.** *Top:* Mouse MCU-variant proteins and the antibody detection sites. *Bottom:* Detection of MCU variant proteins in C2C12 cells overexpressing control vector or shRNA targeted to 3’ UTR of MCU. **F.** *Top, left:* RT-qPCR analysis of MCU-SS using adult FVB mouse tissues. Expression levels are normalized to the heart. *Top, right:* RT-qPCR analysis of MCU-SS using adult cardiomyocytes (ACM) and cardiac fibroblasts (ACF) isolated from adult FVB hearts. Expression levels are normalized to ACM. *Bottom:* Tissue-specific MCU-L/MCU-SS ratio in adult C56BL/6 mice. **G.** PCR of human MCU variants using the total cDNA created from mRNA of HEK293T cells. The fragments were amplified using MCU variant-specific primers and a MCU total primer that reacts with all MCU variants. **H.** RT-qPCR analysis of total MCU and MCU-S in human tissues/cells. Expression levels are normalized to the platelets. **I.** Tissue-specific MCU-L/MCU-S ratio in human platelets and tissues.

In humans, two variants (termed MCU-L and MCU-L2) use the same 5’ exon (exon 1), which encodes the whole component of MTS (Claros & Vincens, 1996) and differs by alternative splicing of exon 4 or 5. Additional levels of complexity in the regulation of MCU may come from the presence of two different gene promoters found in the Eukaryotic Promoter Database (Dreos et al, 2015) (i.e., alternative initiation (Miura et al, 2012))), one of which drives the transcription of MCU-L and -L2 starting from exon 1 (**Fig. 1A** and **Appendix Figure S1**). The order accounts for the transcription of the “short-form” variant (termed MCU-S), which starts with exon 2. MCU- S lacks an MTS since the amino acids of the predicted MTS are not present in exon 2 (**Fig. 1A**, **Appendix Figure S1, and Appendix Table. S1**).

In mice, we also found a candidate transcript that is shorter than human MCU-S (termed MCU-“SS”) (**Fig. 1B and Appendix Figure S1**). By designing specific primers targeting each variant (**Appendix Table S2**), the expression of MCU variants was confirmed in mouse cells and tissues using conventional PCR, quantitative real-time PCR (qPCR), and Western blotting of protein lysates from C2C12 cells stably overexpressing shRNA targeting the 3’ untranslated region (UTR) of CCDC109A (Baughman et al, 2011) (**Fig. 1D-F**). The qPCR assay showed ubiquitous expression of MCU-SS in all investigated tissues/cells with the highest expression on the lung and with various MCU-L/MCU-SS ratios among the different mouse tissues (**Fig. 1F**). We also separately extracted the mRNA from cardiomyocytes and cardiac fibroblasts isolated from adult mouse hearts and found that cardiac fibroblasts have higher expression of MCU-SS compared to cardiomyocytes (**Fig. 1F**).

We next determined the expression of MCU variants in humans. The specific primers targeted for MCU-L2 and MCU-S were designed and their specificity was confirmed by sequencing the PCR products purified from conventional PCR gels (**Appendix Table S2 and Appendix Fig. S2**). The design of specific primers targeting human MCU-L was challenging due to high similarity with other variants (**Appendix Fig. 1)**. Therefore, a primer pair targeting pore region was used to detect all MCU variants (i..e., MCU total), and the MCU-L mRNA amount was then estimated by subtraction of other variants from MCU total mRNA (**Appendix Table S2 and Fig. EV1**). The expression of MCU-L2 was barely detectable by conventional PCR and qPCR in all tissues and cell lines investigated (**Fig. 1G and Fig. EV1**). MCU-S mRNA expression was detected in all investigated cells/tissues, with significantly higher expression in platelets (**Fig. 1G to I, and Fig. EV1**).

In summary, we found N-terminal truncated MCU variants in addition to the originally reported MTS-containing MCU both in human and mouse, we observed particularly high expression of this short variant in human platelets.

### Short MCU variants do not serve as mitochondria channels, but form Ca^2+^ channels at the plasma membrane

Since the first exon for MCU-L which includes MTS is not utilized in human MCU-S or mouse MCU–SS (**Fig. 1 and Appendix Figure S1**), both of them have a much lower probability for mitochondrial transport than MCU-L, as calculated by programs predicting mitochondrial targeting (Almagro Armenteros et al, 2019, Claros & Vincens, 1996) (**Appendix Table S1**). Therefore, we examined whether the MCU-S may be transported into other organelles and/or membrane structures, such as PM, but not to mitochondrial membranes. To test the role of MCU-S in mtCa^2+^ handling, we reintroduced a single MCU variant with a C-terminal tag of GFP or FLAG into MCU knock-down (MCU-KD) HEK293T cell line stably overexpressing shRNA plasmids targeting the 3’-UTR of MCU mRNA (Baughman et al, 2011) (**Appendix Figure S3**).

As reported elsewhere (Baughman et al, 2011, De Stefani et al, 2011, O-Uchi et al, 2013), MCU-L-GFP was exclusively expressed at mitochondria in MCU KO cells, confirmed by co-transfection with mt-RFP (O-Uchi et al, 2013, O-Uchi et al, 2014, Yoon et al, 2003) (**Fig. 2A to C**). In contrast, both MCU-S and MCU-SS showed significantly lower co-localization with mt-RFP, suggesting that these proteins are expressed mainly outside mitochondria including PM (**Fig. 2A-C, and Appendix Figure S3**). MCU-S expression at the PM was also confirmed using a cell surface biotinylation assay (**Fig. 2D**).

**Fig. 2.**
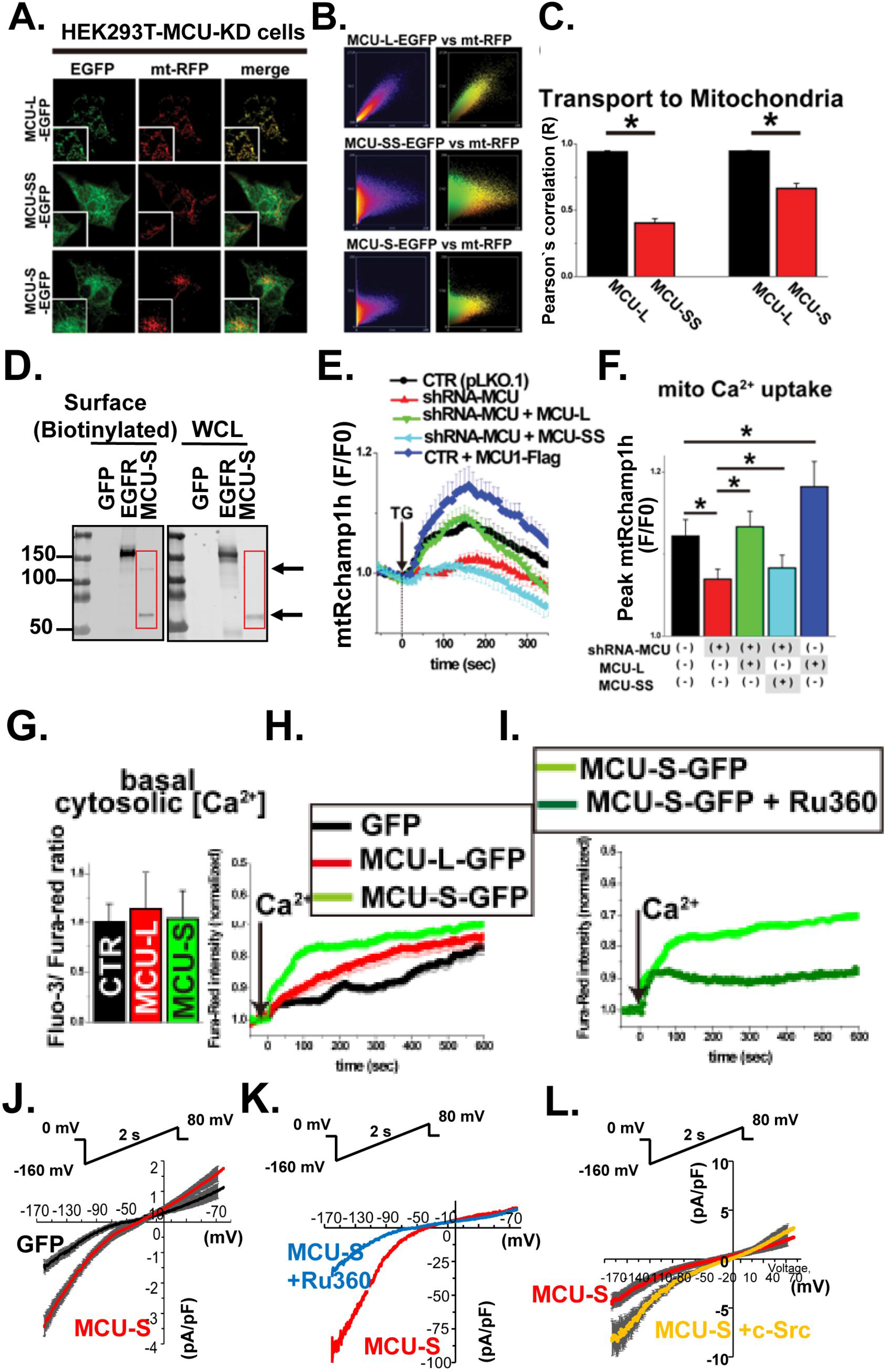
Short MCU variants do not serve as a mitochondrial Ca^2+^ channel, but form Ca^2+^ channels at the plasma membrane. **A.** Subcellular localization of GFP-tagged MCU variants in MCU-KD cells **B.** Frequency (left) and color scatter plots (right) obtained from panel A**. C**. Summary data of Pearson’s correlation coefficient value. The value of Pearson’s correlation coefficient nearly reached 1.0 when when the subcellular localization of GFP and mt-RFP completely overlapped (Jhun et al, 2012, O-Uchi et al, 2013). **D.** Detection of MCU-S- GFP at PM. PM proteins were collected from HEK293Tcells stably overexpressing MCU-S-GFP by cell surface biotinylation. The PM proteins obtained from HEK293T cells stably overexpressing EGFR-GFP were used for positive control. Immunoreactive bands were detected by GFP antibody. In the PM, MCU-S-GFP was detected as monomer and dimer sizes indicated by arrows. **E**. mtCa^2+^ uptake triggered by the SERCA inhibitor 3 μM thapsigargin (TG) in MCU-KD cells after reintroduction of MCU variants. The changes in Ca^2+^ concentration at the mitochondrial matrix [Ca^2+^]_m_ was assessed by mitochondrial matrix-targeted Ca^2+^ biosensor mtRCaMP1h. **F.** Summary data of E. **G.** Basal [Ca^2+^]_c_ in cells stably overexpressing pcDNA (as a control), MCU-L and MCU-S assessed by ratiometric [Ca^2+^] measurement using Ca^2+^-sensitive dye Fluo-3 and Fura-Red. **H.** Changes in [Ca^2+^]_c_ in response to re-admission of 2 mM Ca^2+^ (arrow) following short-time removal of the extracellular Ca^2+^ assessed by Fura-Red. **I.** Effect of 1 μM Ru360 on cells overexpressing MCU-S-EGFP. **J.** Whole-cell *I_ca_* elicited by voltage ramps in HEK293T stably expressing GFP and MCU-S-GFP. **K.** Representative recording of *I_Ca_* before and after 1 μM Ru360 bath application. **L.** Co-expression of constitutively active c-Src enhances whole-cell *I_Ca_* in HEK293T stably expressing MCU- S-GFP.

Reintroduction of MCU-L into MCU-KD cells restored mtCa^2+^ uptake in response to the elevation of cytosolic Ca^2+^ concentration ([Ca^2+^]_c_) by thapsigargin (TG), a sarco/endoplasmic reticulum (ER/SR) Ca^2+^-ATPase (SERCA) inhibitor, close to the levels in control cells, whereas MCU-SS re-expression did not rescue the diminished mtCa^2+^ uptake in MCU-KD cells (**Fig. 2E and F**). To test whether MCU-S forms a functional Ca^2+^-permeable channel at the PM, the changes in [Ca^2+^]_c_ were monitored after re-administration of extracellular Ca^2+^ following its short-time removal. Resting [Ca^2+^]_c_ was not significantly different between MCU-L and MCU-S overexpressing cells (**Fig. 2H**). Upon increase of extracellular Ca^2+^ concentration from 0 to 2 mM, cells stably overexpressing MCU-S-GFP showed [Ca^2+^]_c_ elevation with faster kinetics compared to cells stably overexpressing GFP or MCU-L-GFP, which was significantly inhibited by a 5-min pretreatment with the cell-impermeable MCU inhibitor Ru360 in the extracellular solution (**Fig. 2I**). In contrast, Ru360 showed no significant effect on the [Ca^2+^]_c_ profile in control cells (**Appendix Figure S4**). Next, to precisely characterize mtCUC-like channel function formed by MCU-S at PM, whole-cell Ca^2+^ current (*I_Ca_*) was measured by applying the mtCUC current measurement protocol in whole mitoplast patch (Kirichok et al, 2004). We recorded Ru360-sensitive whole-cell *I_Ca_* elicited by voltage ramps, which is similar to the whole-IMM *mtCUC* current observed in mitoplasts (Kirichok et al, 2004) (**Fig. 2J and K**). Several protein tyrosine kinases (c-Src and Pyk2) activate mtCUC via tyrosine phosphorylation of both MCU termini located in the mitochondrial matrix (Jhun et al, 2016, O-Uchi et al, 2014, Xie et al, 2018). Larger *I_Ca_* was recorded when constitutively active c-Src was co-expressed (**Fig. 2L**), indicating that the N- and C-termini of PM-localized MCU-S face the cytoplasm.

In summary, short MCU variants form Ru360-sensitive Ca^2+^-permeable channels at the PM rather than IMM.

### Short MCU variant inhibits mitochondrial Ca^2+^ uptake

Since MCU-S includes all key structures that are required for maintaining channel activity in MCU-L (Baughman et al, 2011, De Stefani et al, 2011) (see **Fig. 1** and **Appendix Fig. S1**), we next investigated whether MCU-S is capable to interact with MCU-L and affect conventional MCU function. Co-Immunoprecipitation of heterologous co-expression of differentially tagged MCUs (O-Uchi et al, 2014, Raffaello et al, 2013) in HEK293T cells suggests MCU-L - MCU-S interaction in addition to the MCU-L - MCU-L interaction (**Fig. 3A**). Mitochondrial fractionation assay showed that overexpressed MCU-S-GFP in HEK293T cells is exclusively found in the cytosolic fraction (**Fig. EV2**), which contains the ER and PM (O-Uchi et al, 2013, O-Uchi et al, 2014). In MCU-S-GFP (∼55 kDa)-overexpressing HEK293T cells, endogenous MCU-L expression in the mitochondria-enriched fraction (the ∼32-kDa bands in **Fig. EV2A**) was markedly decreased compared to control cells, concomitant with the appearance of a ∼40-kDa band (corresponding to the size of endogenous MCU-L with MTS) in the cytosolic fraction (**Fig. EV2A**). Corresponding to the decreased MCU-L in the mitochondria (**Supplementary Fig. EV2A and B**), MCU-S-GFP overexpression decreased mtCa^2+^ uptake in response to TG application, without affecting the ER Ca^2+^ content or the mitochondrial membrane potential (*Ψ_m_*) (**Fig. 3B to D**). This observation is consistent with overexpression studies of dominant-negative (DN) pore-forming subunits of MCU-L (Baughman et al, 2011, De Stefani et al, 2011, O-Uchi et al, 2014) (MCU-DN) or endogenous MCU-inhibitory subunit MCUb (O-Uchi et al, 2014, Raffaello et al, 2013), which binds to MCU-L and forms Ca^2+^-impermeable pores, but likely via different molecular mechanism (i.e., decreasing the mtCUC number in the mitochondria, see **Fig. EV2C**). Similar results were observed in the primary cells. Overexpressed MCU-S was found outside of mitochondria, inhibits mtCa^2+^ uptake, and inhibits mtCa^2+^- uptake-mediated mitochondrial superoxide generation in rat neonatal cardiomyocytes (**Fig. 3E to G**).

**Fig. 3.**
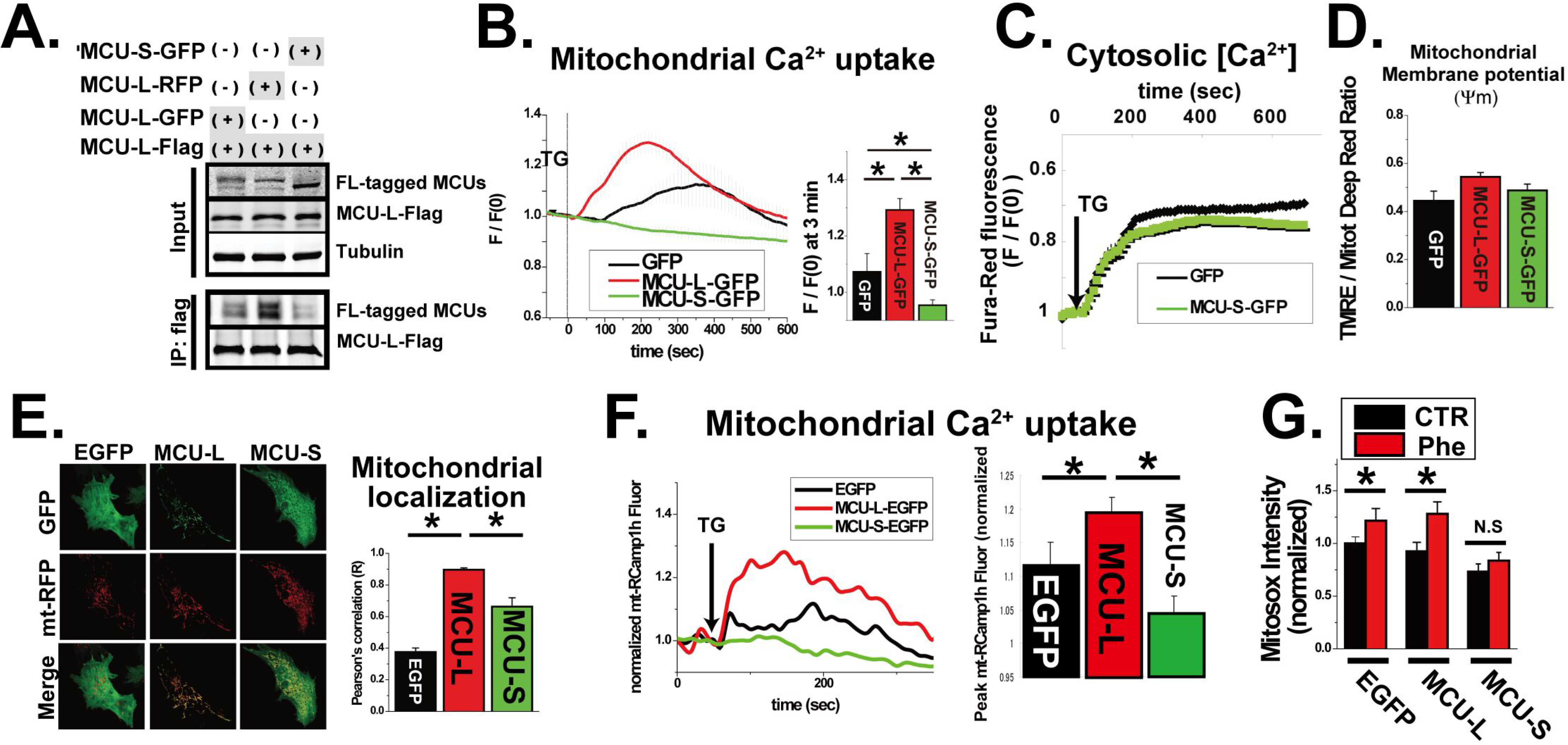
MCU-S inhibits mitochondrial Ca^2+^ uptake in the cardiomyocytes. **A.** Co-immunoprecipitation analysis of MCU-L and MCU-S. HEK293T cells stably overexpressing MCU-L-flag was transiently transfected with fluorescence (FL)-tagged MCU-L or MCU-S and the whole cell lysates were prepared. Co-immunoprecipitation was performed with flag antibody and MCU was detected by antibody A used in Fig. 1E. **B.** mtCa^2+^ uptake triggered by thapsigargin (TG) assessed by mtRCaMP1h. **C.** ER Ca^2+^ estimated by the cytosolic Ca^2+^ elevation levels by TG treatment measured by a Ca^2+^- sensitive dye Fura-Red. **D.** Effect of MCU-S overexpression on resting *Ψ_m_*. Tetramethylrhodamine, ethyl ester (TMRE) is a *Ψ_m_*-sensitive dyes and MitoTracker Deep Red FM (MtDRFM) can label active or inactive mitochondria, which does not depend on *Ψ_m_*. TMRE/MtDRFM ratio was used for comparison. **E.** *Left:* Subcellular localization of GFP-tagged MCU variants in rat neonatal cardiomyocytes (NCMs). EGFP was used as a control. Mitochondria was labeled by mt-RFP. *Right:* Summary data of Pearson’s correlation coefficient values. **F.** *Left:* Representative traces of mtCa^2+^ uptake triggered by TG. [Ca^2+^]_m_ was measured by mtRCaMP1h. *Right:* summary data. \**p*<0.05. **G.** Effect of G_q_-coupled α_1_-adrenoceptor stimulation by phenylephrine (Phe, 100 μM, 0.5 hr) on mitochondrial superoxide levels in NCMs overexpressing MCU variants assessed by MitoSOX-Red. Phe stimulation was used for inducing mitochondrial superoxide generation via mtCa^2+^ overload in NCMs (O-Uchi et al, 2014). NCMs overexpressing MCU-S-GFP have lower mitochondrial superoxide at rest compared to the cells overexpressing EGFP or MCU-L-GFP. NCMs overexpressing MCU-S-GFP did not show the increase of mitochondrial superoxide after Phe stimulation. \**p*<0.05.

In summary, in the presence of MCU-L, MCU-S is capable of forming hetero-oligomers with MCU-L, interfering in MCU-L trafficking to IMM, and subsequently inhibits mtCa^2+^ uptake.

### MCU-S at the plasma membrane forms a Ca^2+^-permeable pathway in human platelets

Since MCU-S mRNA was expressed most abundantly in platelets among the human cells/tissues that we tested (see **Fig. 1**), we addressed next whether endogenous MCU proteins form Ca^2+^ channels at the PM in human platelets from healthy donors, and if so whether this Ca^2+^ pathway has a role in cellular Ca^2+^ signaling. First, three established subunits of mtCUC, MCU, and MICU1/2, (Jhun et al, 2016) were found in the PM proteins collected via a cell surface biotinylation assay (Jhun et al, 2012), but EMRE was not detectable (**Fig. 4A**). Next, the mitochondrial and extra-mitochondrial localization of MCU was confirmed via immunocytochemistry (**Fig. 4B**, see also **Appendix Fig. S5**). Lastly, similar to MCU-S overexpressed HEK293T cells, human platelets exhibited Ru360-sensitive [Ca^2+^]_c_ elevation upon an increase of extracellular [Ca^2+^] from 0 to 3 mM (**Fig. 4C&D**) even in the presence of a specific inhibitor of store-operated calcium entry (SOCE), YM-58483 (Harper & Poole, 2011) (**Fig. 4F&G**). Ru360 did not inhibit SOCE in the platelets (**Fig. 4E&H**).

**Fig. 4.**
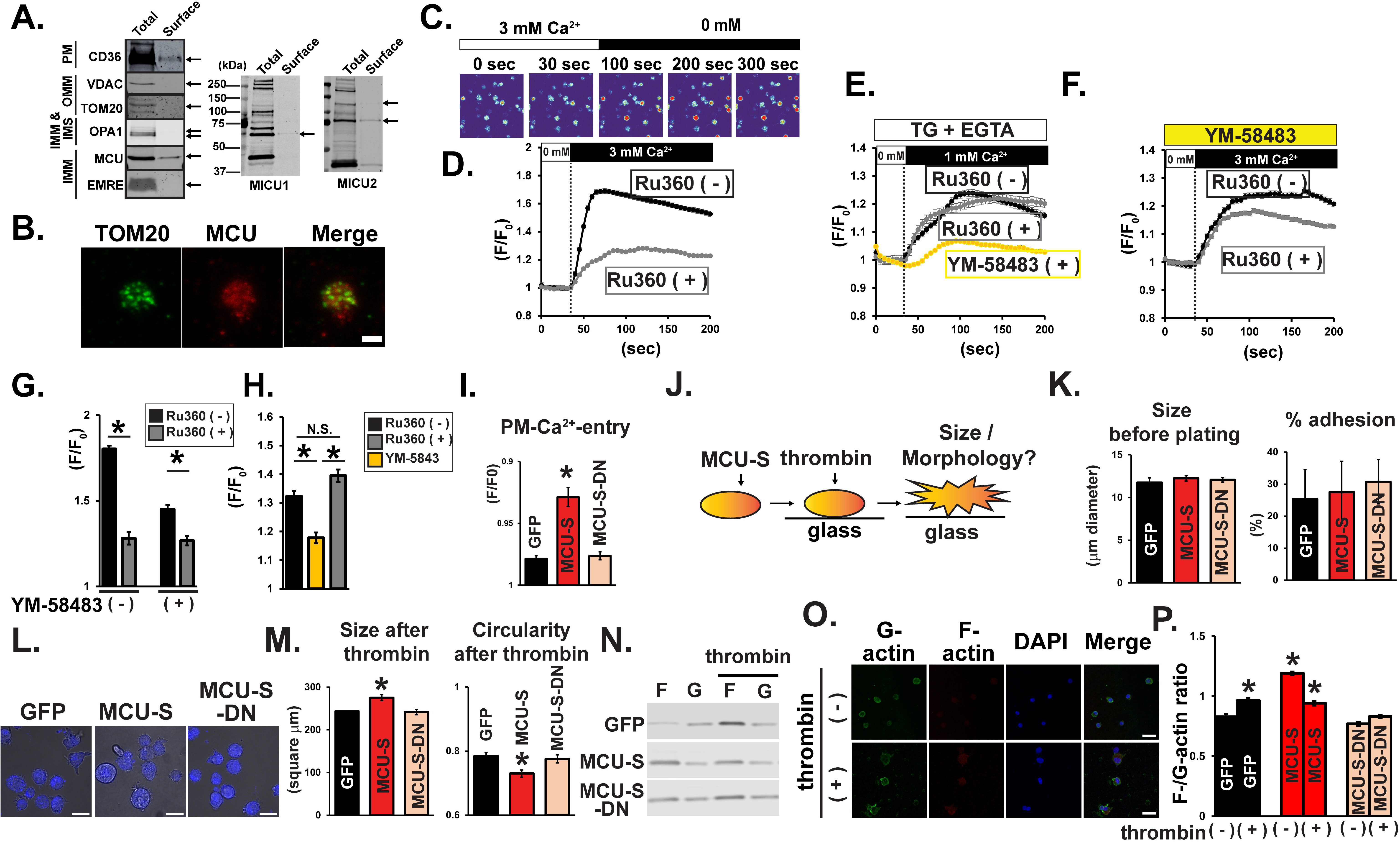
Ca^2+^-permeable MCU channels at the plasma membrane participate in cellular Ca^2+^ signaling in human platelets. **A.** The PM localization of MCU and mtCUC subunits by cell surface biotinylation assay followed by immunoblots. CD36 was used as a positive control for PM proteins in platelets. **B.** Representative confocal images of fixed platelets co-stained with anti-TOM20 and an MCU-specific antibody against the N-terminal domain (see also Supplementary Fig. 7). Scale bar, 2 μm. **C.** Representative time-lapse confocal images of the changes in [Ca^2+^]_cyto_ in response to the reintroduction of extracellular Ca^2+^ (3 mM) after short-term removal assessed by Fluo-3. **D.** Effect of 10 μM Ru360 on Ca^2+^ entry via the PM. **E.** Effect of 10 μM Ru360 and a SOCE inhibitor (3 μM YM-58483) on SOCE. For ER Ca^2+^ depletion, platelets were pretreated with 1 μm TG and 200 μm EGTA for 5 min (Harper & Poole, 2011). **F.** Non-SOCE Ca^2+^ entry via the PM in the presence or absence of Ru360. **G.** Summary data of D&F. \**p*<0.05. **H.** Summary data of E. **I.** Effect of MCU-S or MCU-S-DN expression on Ca^2+^ entry via the PM in MEG-01 cells assessed by Fura-Red. See protocol in panel 4C. \**p*<0.05, compared to GFP-transfected cells. **J.** Schematic diagram of experimental setup. Non-adherent MEG-01 cells were collected, and plated on glass bottom dishes for 20 minutes. Adherent cells were loaded with a cell-permeant fluorescent probe Calcein Violet-AM for 20 minutes to confirm their viability, and then stimulated with thrombin (0.1 U/mL) for 30 minutes. **K.** Effect of MCU-S on cell size (before plating) and adhesion to glass dishes. **L.** Representative confocal and phase contrast images of MEG-01 cells transfected with GFP, MCU-S-GFP, and MCU-S-DN-GFP. Scale bars, 20 μm. **M.** Effect of thrombin on cell size and cell circularity after plating on glass-bottom micro dishes assessed by phase contrast images. Circularity reaches 1 if the cell is perfect circle, and circularity becomes low in complex cell shapes. \**p*<0.05, compared to GFP-transfected cells. **N.** Representative immunoblots of F-/G-actin in MCU-S-GFP transfected MEG-01 cells before and after thrombin stimulation. GFP-and MCU-S-DN-GFP-transfected cells show as comparisons. **O.** Representative confocal images of fixed GFP-transfected MEG-01 cells before and after thrombin stimulation co-stained with monoclonal JLA20 anti-actin antibody and CF568 conjugated-phalloidin to detect G-actin and F-actin, respectively. Scale bars, Scale bars, 20 μm. **P.** Summary data of F-/G-actin ratio quantified from confocal images. \**p*<0.05, compared to GFP-transfected cells before thrombin stimulation.

In summary, the human platelets possess an endogenous Ru360-sensitve Ca^2+^- permeable pathway at the PM that is an independent mechanism from SOCE.

Since *in vitro* gene modification is challenging in platelets due to lacking their nucleus, we next utilized a human megakaryocyte leukemia cell line (MEG-01) to further assess the impact of PM-MCU channel function to the cellular Ca^2+^ signaling. Importantly, MCU-S mRNA was barely detectable in MEG-01 cells and the conventional protocols for the induction of *in vitro* differentiation/maturation (Dhenge et al, 2019), which leads to polyploidy and the production of platelet-like particles, failed to induce the MCU-S expression in MEG-01 cells, indicating that MCU-S expression is likely specific to the matured platelets *in vivo* (**Appendix Fig. S6**). Therefore, we used this cell line as a template for exogenously expressing MCU-S and its dominant-negative (DN) mutant MCU-S-DN (mutating negatively charged residues of the conserved region in the MCU-S pore [D260, E263 in MCU-L (O-Uchi et al, 2014)] into glutamines). Introducing MCU-S, but not MCU-S-DN, increased PM-Ca^2+^ entry, as we observed in the HEK293T system (**Fig. 4I**). Platelets undergo morphologic changes via actin polymerization when [Ca^2+^]_c_ elevation occurs such as ER Ca^2+^ release by thrombin stimulation, one of the key inducers of platelet activation *in vivo* via protease-activated receptors (PARs) (Bearer et al, 2002). Following the published protocol (Heo et al, 2022), we next plated non-adherent MEG-01 cells to adhere to glass bottom microplates, and then treated with thrombin to activate downstream Ca^2+^ signaling (**Fig. 4J**). MCU-S or MCU-S-DN expression did not affect the cell size before plating, adhesion efficiency (**Fig. 4K**), or the expression of cellular Ca^2+^ handling proteins including SOCE machinery proteins, ER Ca^2+^ handling proteins, mtCa^2+^ handling proteins, and conventional PM-Ca^2+^ entry pathways in platelets (**Fig. EV3)**. First, morphological changes by thrombin treatment in MEG-01 cells were not affected by the pretreatment of a mitochondrial uncoupler FCCP that can eliminate the mtCa^2+^ uptake (**Fig. EV4**). This indicates that 1) the Ca^2+^ release via mitochondrial permeability transition pore (mPTP) triggered by the mtCa^2+^ uptake is likely not required for actin activation and 2) [Ca^2+^]_c_ elevation by ER Ca^2+^ release is sufficient to cause morphologic changes. Importantly, MCU-S transfected cells showed increased cell size and decreased circularity after thrombin stimulation compared to GFP (as a control) and MCU-S-DN transfected cells (**Fig. 4L&M**). Moreover, this effect by MCU-S was enhanced by the FCCP pretreatment possibly due to the decreased [Ca^2+^]_c_ buffering by mitochondria that exaggerates the [Ca^2+^]_c_ elevation (**Fig. EV4**). Lastly, we confirmed that thrombin increased β-actin polymerization (as assessed by increased filamentous [F]-actin /globular [G]-actin ratio (Antonipillai et al, 2019, Dasgupta et al, 2016, Lee et al, 2013))) in control cells, whereas in MCU-S-transfected cells, F-/G-actin ratio was already elevated without thrombin stimulation. Lastly, thrombin-induced β-actin polymerization was abolished by MCU-S-DN transfection (**Fig. 4N-P**).

In summary, endogenous MCU proteins, likely MCU-S, form Ca^2+^ channels at the PM, the Ca^2+^ entry via PM-MCU-S channels activate Ca^2+^ signaling for actin rearrangements after global [Ca^2+^]_c_ elevation by thrombin, which is critical for the activation of aggregation/coagulation pathways in platelets (Abbasian et al, 2020, Denorme & Campbell, 2022, Millington-Burgess & Harper, 2021).

## Discussion

MCU is dogmatically known as a pore subunit for the highly Ca^2+^-selective channel at the IMM that represents the major mechanism to take up mtCa^2+^. This is mainly because 1) the MTS found in the conventional MCU-L variant and 2) mass spectrometry data from human and mouse mitochondria, such as MitoCarta (Calvo et al, 2016), strongly support MCU-L protein expression in the mitochondria. Here, we reported that MCU-S variant that does not possess an MTS (**Fig. 1**) enables the MCU tetramers outside of mitochondria (**Fig. 3**) and is capable of forming mtCUC-like Ca^2+^-permeable channels at the PM in a heterologous expression system and in human platelets (**Figs. 2&4**).

The human mtCUC contains 4 or 5 major subunits: 1) MCU which tetramerizes to form the Ca^2+^-permeable pore for mtCUC; 2) EMRE, a single-pass transmembrane protein that activates metazoan MCU channel activity in the IMM environment; and 3) MICU family members that provide Ca^2+^-sensitivity to the mtCUC. In the PM of human platelets, we detected MCU, MICU1 and 2, but not EMRE (**Fig. 4A**). While there is an important role of EMRE in Ca^2+^ transport activity via MCU oligomers in the IMM in higher organisms, it has been shown that the human MCU protein alone is sufficient for MCU activity in artificial bilayers (De Stefani et al, 2011, Patron et al, 2014, Raffaello et al, 2013). Since the lipid composition/environment is different between the IMM and PM, MCU-S oligomers may produce Ca^2+^ permeability without interacting with EMRE in the PM, similar to the case in artificial bilayers. Interaction between the MICU complex with MCU is also suggested via EMRE, but the recombinant MICU1 and 2 are capable of manipulating the channel activity of human MCU protein oligomers in artificial bilayers without EMRE (Patron et al, 2014). This observation also suggests that the MICU1- MICU2 complex might regulate MCU-S channels at the PM in the absence of MCU-EMRE association. Nevertheless, given that EMRE exerts significant influence on intrinsic channel activity, and/or channel complex assembly in the IMM, the detailed channel properties of mtCUC at the IMM and MCU-S channels at the PM, such as single channel conductance and the Ca^2+^-sensitivity, are likely different. However, our data using MCU-S-DN clearly provides evidence that MCU channels at the PM have a similar putative pore-forming structure that produces Ca^2+^ permeability of mtCUC.

Our MCU-S overexpression studies showed that MCU-L/MCU-S expression ratios regulate the subcellular localization of MCU tetramers (**Fig. 3 and Fig. EV2**), suggesting that the majority of human cell types/tissues (see **Fig. 1**) we tested may not possess endogenous MCU channels outside of mitochondria except platelets. Indeed, we detected a Ru360-sensitive Ca^2+^-permeable pathway at the PM, which is independent from SOCE in the human platelets (**Fig. 4**). Human platelets have three major molecular mechanisms that elevate [Ca^2+^]_c_ during their activation (Millington-Burgess & Harper, 2021, Varga-Szabo et al, 2009): 1) ER Ca^2+^ release; 2) Ca^2+^ entry via PM channels including SOCE, transient receptor potential channel family, and receptor-operated Ca^2+^ channels; and 3) mPTP opening triggered by the mtCa^2+^ uptake via mtCUC. Supramaximal [Ca^2+^]_c_ signal is required for transformation into procoagulant platelets *in vitro* experiments (Abbasian et al, 2020). Although PM-mediated Ca^2+^ entry mechanism holds most smallest impact on [Ca^2+^]_c_, the assessment of recent studies using transgenic mouse models with knock out of major SOCE components revealed that PM-Ca^2+^ entry mechanism via SOCE is also essential for procoagulant platelet formation (Mammadova-Bach et al, 2019, Millington-Burgess & Harper, 2021, Varga-Szabo et al, 2009). Lastly, we also showed that Ca^2+^ permeability via MCU-S channels at the PM is capable of modulating thrombin-induced actin cytoskeleton remodeling and cell morphological changes (**Fig. 4**), which precede platelet aggregation or adhesion. Recently, the importance of mtCa^2+^ influx and mtCUC-mediated mPTP opening for platelet activation has been shown (Abbasian et al, 2020, Kholmukhamedov et al, 2018). Our observation strongly suggests that the MCU channels expressed outside of the mitochondria are also additionally involved in the platelet activation.

In conclusion, we identified novel MCU variants that lack the MTS, which enables the MCU channels to localize outside of mitochondria. This is completely distinct from the originally reported MCU function as a highly mitochondria-specific Ca^2+^ channel. Expression ratios of the conventional mitochondria-targeted MCU and the non-mitochondria-targeted variants regulate the subcellular localization of the MCU channels. Human MCU variant MCU-S forms Ca^2+^-permeable channels at the PM of platelets and participates in the Ca^2+^- signaling for platelet activation. Our finding provides novel insights into the molecular basis of MCU variant-dependent cellular Ca^2+^ handling in physiological and potentially in pathological conditions in which both cytosolic and mitochondrial Ca^2+^ dynamics play key roles. Since some members of bacteria and fungi groups contain putative MCU homologs without MTS, characterization of MCU variants may also help to elucidate the role of mtCa^2+^ transport in the evolution of mitochondria as well as the emergence of eukaryotic life.

## Materials and Methods

### Ethical approval

All animal experiments were performed in accordance with the Guidelines on Animal Experimentation of Thomas Jefferson University, Rhode Island Hospital, and University of Minnesota. The study protocols were approved by the Institutional Animal Care and Use Committee at each institution. The investigation conformed to the Guidelines for the Care and Use of Laboratory Animals published by the US National Institutes of Health (NIH).

### Antibodies, plasmids, and reagents

The antibodies and plasmids used for the experiments are listed in **Appendix Table. S4 and S5**, respectively. All chemicals and reagents were purchased from Sigma-Aldrich Corporation (St Louis, MO, USA) otherwise indicated; thapsigargin (Alomone lab, Jerusalem, Israel); Ru360 (MilliporeSigma, Burlington, MA); YM-58493 (Selleck Chemicals, Houston, TX); valproic acid (AmBeed, Arlington Heights, IL); (trifluoromethoxy)phenylhydrazone (FCCP) (Focus Biomolecules, Plymouth Meeting, PA); MitoSOX-Red and MitoTracker Deep Red FM (mtDRFM) (Thermo Fisher Scientific, Waltham, MA); and Tetramethylrhodamine ethyl ester perchlorate (TMRE) (Biotium, Fremont, CA).

### Variants information acquisition and sequence analysis

The human and mouse MCU gene and variants information were sourced from the Ensemble database (https://useast.ensembl.org/). Conserved domains of variants were analyzed by the Clustal Omega multiple sequence alignment program of the European Bioinformatics Institute (http://www.ebi.ac.uk). The mitochondria targeting probabilities of variants were determined using the MitoProt II prediction program (Claros & Vincens, 1996).

### RNA extraction, reverse transcription, and quantitative Real-Time PCR (qRT-PCR)

Total RNA was isolated from the mouse cells/tissues and human cell lines using the RNeasy® Mini Kit (Qiagen Inc., Germantown, MD). For human platelets and MEG-01 cells, total RNA was isolated using TRIzol™ Reagent (Therrmo Fisher Scientific, Waltham, MA). RNA quantity and purity were measured using a NanoDrop 2000 spectrophotometer (NanoDrop Technologies, Wilmington, DE). cDNA was synthesized from 1.0 or 2.0 µg of total RNA using the iScript™ reverse transcription (Bio-Rad Laboratories, Hercules, CA) kit or Superscript™ IV VILO™ Master Mix (Thermo Fisher Scientific), respectively. Total cDNA from various human tissues were purchased from Biochain (Newark, CA), Takara Bio USA (Ann Arbor, MI), and Zyagen (San Diego, CA). Donor information of human total cDNA provided from the companies was listed in **Appendix Table. S6**. The primer set for the amplification of MCU cDNA was designed according to Ensembl sequences using the Primer3 program (Untergasser et al, 2012) (**Appendix Table. S2**). All the other primer sets for the assessment for human and mouse cells/tissues were listed in **Appendix Table. S3**. All the primers were purchased from Integrated DNA Technologies, Inc. (Coralville, IA) and Sigma-Aldrich. The PCR products of human MCU variants from HEK293T cells were separated by electrophoresis on 2.0% agarose gels in Tris-borate-EDTA (TBE) buffer, stained with ethidium bromide (Thermo Fisher Scientific), and visualized under ultraviolet light. For DNA sequence analysis, amplified PCR products were extracted from agarose gel using the QIAquick gel extraction kit (Qiagen) and were sequenced by the dideoxy (Sanger) approach at Integrated DNA Technologies, Inc. Sequences were further compared with the full-length human MCU mRNA sequence (**Appendix Fig. S2)**.

qRT-PCR for mouse cells/tissues, and human cell lines in Fig. 1F and Supplementary Fig. 3 was performed using an iCycler thermocycler (Bio-Rad laboratories) with a TaqMan RT-PCR Master Mix (Bio-Rad Laboratories) or Fast SYBR Green Master Mix (Thermo Fisher Scientific) with Applied Biosystems 7500 Fast (Thermo Fisher Scientific).

qRT-PCR for human platelets and tissues in Fig. 1H was performed using an Applied Biosystems 7500 Fast (Thermo Fisher Scientific) with an Applied Biosystems PowerUp SYBR Green Master Mix (Thermo Fisher Scientific). qRT-PCR for MEG-01 cells in Supplementary Fig. 9 was performed using a QuantStudio™ 3 (Thermo Fisher Scientific) with an Applied Biosystems PowerUp SYBR Green Master Mix (Thermo Fisher Scientific). The relative transcripts level for human and mouse MCU variants were determined by the ΔΔC_T_ method with GAPDH as the endogenous reference gene.

### Cells, Culture, and transfection

All the cell lines were purchased from American Type Culture Collection (Manassas, VA) otherwise indicated. Human platelets from healthy donors were purchased from Zen-Bio (Durham, NC) and HumanCells Biosciences (Fremont, CA). Donor information provided from the companies was listed in **Appendix Table. S7**.

HEK293T cells (kindly provided by Dr Keigi Fujiwara) were maintained in Dulbecco’s modified Eagle’s medium (DMEM) (Cytiva, Marlborough, MA) supplemented with 4.5 g/L glucose, 1 mM sodium pyruvate and 1% L-glutamine, 10% fetal bovine serum (FBS) (GIBCO, Grand Island, NY), 100 U/ml penicillin, 100 μg/ml streptomycin (Genesee Scientific, San Diego, CA, USA) at 37°C with 5% CO_2_ in a humidified incubator. For transient transfection, HEK293T cells were transfected using FUGENE-HD (Promega, Madison, WI) and used for experiments after 48 to 72 h after transfection as we previously demonstrated (O-Uchi et al, 2014).

C2C12 mouse myoblast cells (kindly provided by Dr. Robert T Dirksen) were maintained in DMEM (Sigma-Aldrich) supplemented with 1.0 g/L glucose, 1 mM sodium pyruvate and 1% L-glutamine, 20% FBS (GIBCO), 100 U/ml penicillin, 100 μg/ml streptomycin and 0.25 μg/mL Fungizone (GIBCO) and at 37°C with 5% CO_2_ in a humidified incubator.

MEG-01 cells were maintained in RPMI 1640 medium (Cytiva and Genesee Scientific) supplemented with 10% fetal bovine serum (FBS) (GIBCO), 100 U/ml penicillin, 100 μg/ml streptomycin (Genesee Scientific) at 37°C with 5% CO_2_ in a humidified incubator. For transient transfection, cells were transfected using FUGENE-HD (Promega) and used for experiments after 48 to 72 h after transfection. Cell counting and cell size measurements before and after the adhesion on the glass bottom plates were performed by EVE™ Automated Cell Counter, NanoEnTek (Seoul, Republic of Korea).

HEK293T cells stably overexpressing MCU variants, and EGFR-GFP were generated as we previously reported (O-Uchi et al, 2014, Vang et al, 2021). Cells stably overexpressing pcDNA3.1(+) or pEGFP-N1 were used as controls. HEK293T and C2C12 cells stably knocking down MCU were generated by transfecting with plasmids carrying shRNA targeted 3’UTR of human and mouse MCU, respectively. Cells stably overexpressing empty PLKO.1 plasmid was used as a control. Stable cell lines were maintained in 1200 μg/ml G-418 (Mediatech/Corning, Corning NY) or 0.5μg /ml puromycin (Gemini Bio Products, West Sacramento, CA).

Neonatal cardiomyocytes (NCMs) were enzymatically isolated from ventricles from two-to three-day-old Sprague Dawley rats (Envigo, Huntingdon, UK and Taconic Biosciences, Germantown, NY) using collagenase type 2 (Worthington Biochemical Corp., Lakewood, NJ, USA) as we previously reported (Jhun et al, 2018). Rats were euthanized by decapitation and hearts were harvested for NCM isolation. NCMs were plated onto gelatin-coated dishes and maintained in DMEM (Mediatech/Corning) supplemented with 4.5 g/L glucose, 1 mM sodium pyruvate and 1% L-glutamine, 10% FBS (GIBCO), 100 U/ml penicillin, 10 mg/ml streptomycin (Mediatech/Corning) s at 37°C with 5% CO_2_ in a humidified incubator^3^. NCMs were transfected using FUGENE-HD (Promega) 48 h after isolation and used for experiments after 48 h after transfection (O-Uchi et al, 2014).

Adult cardiomyocytes (ACMs) and cardiac fibroblasts (ACFs) were isolated from ventricles of 8-month-old adult FVB hearts (Taconic, Germantown, NY) by retrograde heart perfusion with collagenase type II (Worthington Biochemical Corp). ACM were allowed to settle by gravity in the washing buffer with 2% BSA and stepwise [Ca^2+^] increase from 20 μM to 1.0 mM, and then used fresh. ACF were cultured in DMEM/F12 (Thermo Fisher Scientific) with 10% FBS, 100 U/ml penicillin, and 100 mg/ml streptomycin, and collected after 48 hrs.

### Protein fractionations, Western blot analysis, immunoprecipitation

Whole cell lysates were prepared using commercial lysis buffer (Cell Signaling Technology, Danvers, MA) with 1 mM PMSF and 1% protease inhibitor cocktail (Sigma-Aldrich). Mitochondria-enriched protein fraction and cytosolic protein fraction containing endoplasmic reticulum proteins and plasma-membrane proteins were isolated as previously described with different centrifugation (Jhun et al, 2018, O-Uchi et al, 2014).

Immunoprecipitation was performed as we previously described (Jhun et al, 2018, O-Uchi et al, 2014). Briefly, protein lysates (500 μg) were incubated with primary antibodies (1 μg) for overnight at 4°C, followed by the addition of Protein A/G Plus Agarose (Santa Cruz Biotechnology, Dallas, TX). Immunocomplexes were washed with lysis buffer and subsequently subjected to Western blot.

For biochemically measuring F-actin/G-actin ratio, Adhered MEG-01 on the glass bottom dishes were detached, collected, washed with PBS, and lysed with actin stabilization buffer containing 100 mM PIPES in pH 6.9, 30% glycerol, 5% DMSO, 1 mM MgSO4, 1 mM EGTA, 1% Triton X-100, 1 mM ATP and 1% protease inhibitor cocktail (Sigma-Aldrich) (Antonipillai et al, 2019, Antonipillai et al, 2020). Lysates were incubated on ice for 10 min and centrifuged at 100,000 x g for 60 min at 4°C. Supernatant was corrected as a fraction containing G-actin. The pellets containing F-actin were solubilized with actin depolymerization buffer containing 100 mM PIPES in pH 6.9, 1 mM MgSO_4_, 10 mM CaCl_2_ and 5 μM cytochalasin D (Focus Biomolecules) and incubated on ice for 30 min (Antonipillai et al, 2019, Antonipillai et al, 2020).

All the protein samples were separated by sodium dodecyl sulfate polyacrylamide (SDS)-PAGE and were transferred to nitrocellulose membrane (Bio-Rad laboratories, Genesee Scientific, Cytiva, and Santa Cruz Biotechnology) and incubated with primary antibodies, followed by the treatment of fluorescence-conjugated secondary antibody (LI-COR Biotechnology, Lincoln, NE). Immunoreactive bands were visualized by Odyssey Infrared Imaging System (LI-COR Biotechnology) and quantified by densitometry using Image J software (NIH) and Image Studio Lite Version 5.2 (LI-COR Biotechnology).

### Cell surface protein biotinylation assay

To distinguish the cell surface localization of MCU, a cell-surface biotinylation assay was performed using Pierce Cell Surface Protein Isolation Kit (Thermo Fisher Scientific).

HEK293T cells stably overexpressing GFP-tagged proteins plated in 15-cm dishes were washed with an ice-cold Ca^2+^ and Mg^2+^-free phosphate-buffered saline (PBS), and labled with EZ-Link Sulfo-NHS-SS-Biotin™ (Thermo Fisher Scientific) for 30 minutes at 4°C, followed by quenching, and harvesting with tris-buffered saline (TBS). For collecting surface protein from human platelets, platelets were rinsed once with PBS at room temperature, and incubated with EZ-Link Sulfo-NHS-SS-Biotin™ (Thermo Fisher Scientific) in PBS for 30 minutes at room temperature. The biotin labeling reaction was quenched with cold PBS containing 100 mM of glycine for 10 minutes.

After labeling, cells and platelets were centrifuged at 500 x g for 3 min at 4°C, and the pellets were treated with lysis buffer containing protease inhibitor cocktail (Sigma-Aldrich), followed by a brief sonication, and incubated on ice for 30 minutes with vortexing every 5 minutes for 5 seconds. Next, cell lysates were centrifuged at 10,000 *× g* for 2 minutes at 4°C, and supernatants were collected in the separated tubes. To isolate labeled proteins supernatants were incubated with NeutrAvidin™ agarose (Themo Fisher Scientific) for 60 minutes at room temperature. Biotinylated protein-NeutrAvidin™ agarose complex was collected, washed by the washing buffer containing protease inhibitor cocktail (Sigma-Aldrich), and eluted by sample buffer (Pierce™ Lane Marker Non-Reducing Sample Buffer, Thermo Fisher Scientific) with 10 mM dithiothreitol (DTT) for 60 minutes at room temperature, followed by a centrifugation at 1,000 *× g* for 2 minutes. Eluted protein samples were mixed with sample buffer and subjected to SDS-PAGE and Western blot.

### Live cell Ca^2+^ imaging

HEK293T cells and NCMs were plated on glass bottom dishes (MatTek, Ashland, MA and Matsunami Glass, Osaka, Japan), loaded with membrane-permeant AM ester forms of Fluo-3 (Fluo-3-AM) (Biotium) and/or Fura-Red, Fura 2-TH-AM (Fura-Red-AM) (Setareh Biotech, Eugene, OR) at 37°C, and the changes in cytosolic Ca^2+^ concentration ([Ca^2+^]_c_) were measured at room temperature by FV1000 laser scanning confocal microscope (Olympus, Tokyo, Japan) in modified HEPES Tyrode’s solution (in mM): NaCl, 136.9; KCl, 5.4; CaCl_2_, 1; MgCl_2_, 0.5; NaH_2_PO_4_, 0.33; HEPES, 5; glucose, 5; pH 7.40 adjusted with NaOH (O-Uchi et al, 2013, O-Uchi et al, 2014, Vang et al, 2021). For measuring the changes in Ca^2+^ concentration at mitochondrial matrix ([Ca^2+^]_mt_), cells were transiently transfected with mitochondrial matrix-targeted Ca^2+^ biosensor mt-RChamp1h and the mt-RChamp1h fluorescence was monitored with excitation wavelength at 543 nm and emission wavelength at 560-660 nm at room temperature (Hamilton et al, 2018). All data were analyzed using Image J (NIH).

Human platelets were resuspended in modified platelet HEPES Tyrode’s buffer (in mM: 136.5 mM NaCl, 136.5; KCl, 2.68; NaHCO_3_, 11.9, NaH_2_PO_4_, 0.42; MgCl_2_, 0.5; HEPES, 5; pH 7.40 adjusted with NaOH) containing 0.35% (w/v) human serum albumin (HSA) 0.35% (w/v), and prostaglandin I2 (PGI2, 0.5 μM) (Cayman Chemical, Ann Arbor, Michigan). Platelets were immobilized on the glass bottom plates (Cellvis Mountain View, CA) coated with Cell-Tak (Corning)-coated, loaded with Fluo-3-AM (Biotium) for 30 min at room temperature, and wash with modified platelet HEPES Tyrode’s buffer containing 0.35% HSA and 0.5 μM PGI_2_. Ca^2+^ imaging was performed in modified platelet HEPES Tyrode’s buffer containing 0.1 % HSA and apyrase (0.02 U/mL) (Harper & Poole, 2011) using an FV3000 laser scanning confocal microscopes (Olympus) at room temperature All data were analyzed using Fiji (Schindelin et al, 2012).

### Quantitative co-localization and morphology analysis in live cells

Colocalization analysis of the live cell images and the generation of color and frequency scatter plots were performed using Image J software (NIH) with an Intensity Correlation Analysis plugin (The Bob and Joan Wright Cell Imaging Facility, Toronto Western Hospital, Toronto, Ontario). Colocalization was assessed using Pearson’s correlation coefficient (Jhun et al, 2018, O-Uchi et al, 2013). The values for Pearson’s correlation range from 1 to −1. A value of 1 represents perfect correlation, −1 represents perfect exclusion, and 0 represents random localization.

For the morphology assay of MEG-01 cells, non-adherent MEG-01 cells were resuspended in modified platelet HEPES Tyrode’s buffer with 0.4 mM CaCl_2_, plated on glass bottom dishes (MatTek). Adherent cells were loaded with a cell-permeant fluorescent probe, Calcein Violet-AM (Biolegend, San Diego, CA), for 20 minutes, followed by the stimulation with thrombin (0.1 U/mL) for 30 min (Heo et al, 2022). Cell morphology was measured under an FV3000 laser scanning confocal microscopes (Olympus) at room temperature. Cells stained with Calcein Violet were defied as viable cells, and their circularity was assessed from phase contrast images using Fiji (Schindelin et al, 2012).

### Immunocytochemistry

Immunocytochemistry for HEK293T cells and human platelets were performed as we previously described (Jhun et al, 2018). Briefly, HEK293T cells plated on glass bottom dishes and platelets immobilized on poly-L-lysine (Electron Microscopy Sciences, Hatfield, PA)-coated on glass bottom dishes were fixed with 4.0 % paraformaldehyde (Thermo Fisher Scientific) at 4°C for 10 min, permeabilized with saponin at room temperature for 10 min, blocked with 10% goat serum (Cell Signaling Technology), and incubated with primary antibodies overnight, followed by incubation with fluorescent secondary antibodies (Thermo Fisher Scientific) for 1 hr.

Immunostaining of F-and G-actin in MEG-01 cells were performed following the published protocol (Lee et al, 2013). Briefly, cells were first fixed with 4.0 % paraformaldehyde (Thermo Fisher Scientific) for 15 min on ice, and washed by cold PBS, followed by additional fixation by cold acetone (Neta Scientific, Hainesport, NJ) at -20°C for 5 min. For detecting G-actin, cells were washed again with cold PBS after fixation, blocked with 10% goat serum (Cell Signaling Technology), and incubated with a monoclonal anti-actin JLA20 antibody (provided by Developmental Studies Hybridoma Bank [DSHB] at University of Iowa, Iowa City. Iowa. JLA20 was deposited to the DSHB by Lin, J.J.-C.) for overnight, followed by incubation with Alexa488-conjugated secondary antibody (Thermo Fisher Scientific) for 1 hr. For staining F-actin, cells were incubated with CF568-conjugated phalloidin (Biotium) and DAPI (Biotium) before observation.

Immuno-stained images were acquired with a an FV3000 laser scanning confocal microscopes (Olympus) at room temperature. Negative control experiments were performed using secondary antibodies without incubation of primary antibodies, which showed no noticeable labelling.

### Electrophysiological measurements

Whole-cell patch-clamp MCU calcium current (*I_Ca, MCU_*) was carried out for measuring the Ca^2+^ current via hMCU-S (*I_Ca, MCU_*) from in HEK293T cells stably overexpressing hMCU-S-GFP at room temperature using an Axopatch 200 amplifier with Digidata 1550B, CV-203BU head stage, and PClamp 10 software (Molecular Devices, San Jose,CA). Briefly, before recordings, culture medium was removed, and cells were washed with Tyrode’s solution containing (in mM): 135, NaCl; 5, KCl; 1, MgCl_2_; 2, CaCl_2_; 10, HEPES; 5.6 Glucose; pH was adjusted to 7.4 with NaOH. For eliminating endogenous K^+^ currents, Tyrode’s solution was modified by substituting KCl with CsCl, and use as a bath solution (in mM): 135, NaCl; 5, CsCl; 1, MgCl_2_; 2, CaCl_2_; 10, HEPES; 5.6, Glucose; pH was adjusted to 7.4 with CsOH. The composition of the pipette solution was set up as nominally free Ca^2+^ as follows (in mM): 135, CsCl; 5, NaCl; 1, MgCl_2_; 10, HEPES; pH was adjusted to 7.2 with CsOH. The *I_Ca, MCU_* were elicited with a standard voltage ramp protocol (Chaudhuri et al, 2013, Kirichok et al, 2004). From a holding potential of 0 mV, a 100-ms step hyperpolarization to -160 mV was followed by an ascending voltage ramp (from -160 mV to +80 mV at 120 mV/s), applied every 5 seconds. MCU blocker Ru360 (1 μM) was added and defined Ru360-sensitive component as *I_Ca, MCU_*. Recording electrodes were pulled from thin-wall borosilicate glass capillaries with filament (World Precision Instruments, Sarasota, FL, USA) using pipette puller (model P-97, Sutter Instruments, Novato, CA, USA). Data were low-pass filtered at 5 kHz and digitized at sampling rate 20 kHz.

### Statistical analysis

All results are shown as mean standard error. Unpaired Student’s t-test was performed for two data sets. For multiple comparisons, one-way ANOVA followed by the post hoc Tukey test was performed. Statistical significance was set as a p value of <0.05.

## Acknowledgments

We thank Ms. Donquin Yang, Ms. Michelle King, Mr. Jocob Mollar, Ms. Jessica Cao (Rhode Island Hospital and Brown University), Dr. Bing Yi, Ms. Sarah Monaco (Thomas Jefferson University), and Dr. Yuta Suzuki (University of Minnesota) for their technical assistance. This The work was supported by NIH/NHLBI R01HL160699 (to B.S.J), NIH NIH/NIGMS U54GM115677 (to B.S.J), American Heart Association, 18CDA34110091 (to B.S.J), NIH/NHLBI R01HL136757 (to J.O.-U.), NIH/NIGMS P30GM1114750 (to J.O.- U.), W.W. Smith Charitable Trust No. H1403 Medical Research Award (to J.O.-U.), Rhode Island Foundation No. 20164376 Medical Research Grant (to J.O.-U.), and American Physiological Society, 2017 Shih-Chun Wang Young Investigator Award (to J.O.-U.).

## Expanded View Figure legends

**Fig. EV1.**
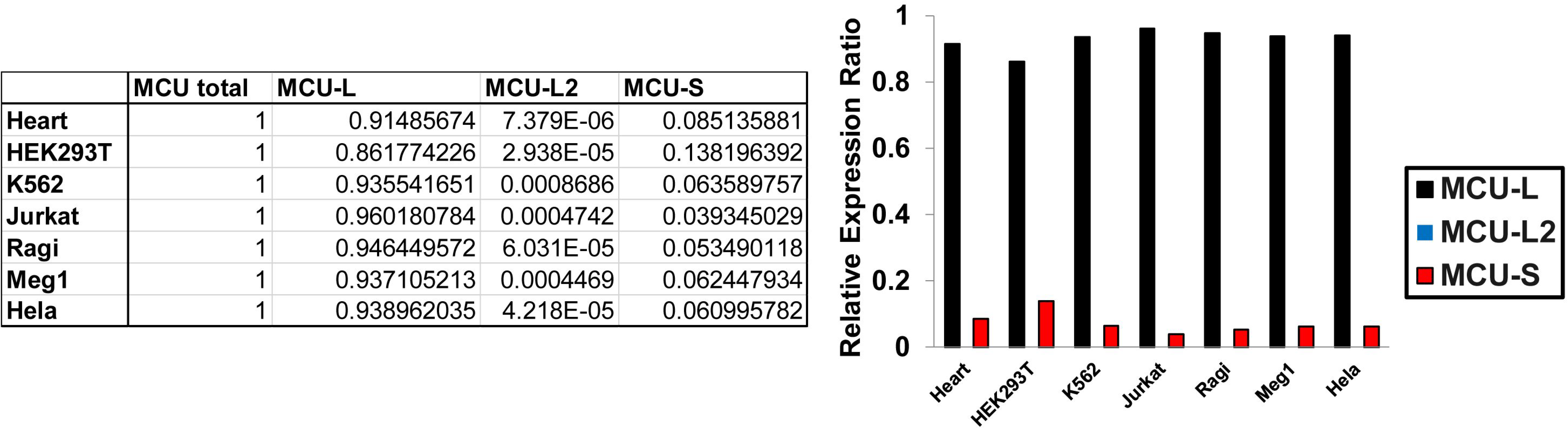
Real time qPCR analysis of MCU variants in human cell lines. Human MCU-L2 and MUC-S were measured using specific primers (see **Supplementary Fig. 2** and **Supplementary Table 2**). The total MCU primer can detect all MCU variants (**Supplementary Table 2**). Representative results are shown in the table (left) and the bar graph (right).

**Fig. EV2.**
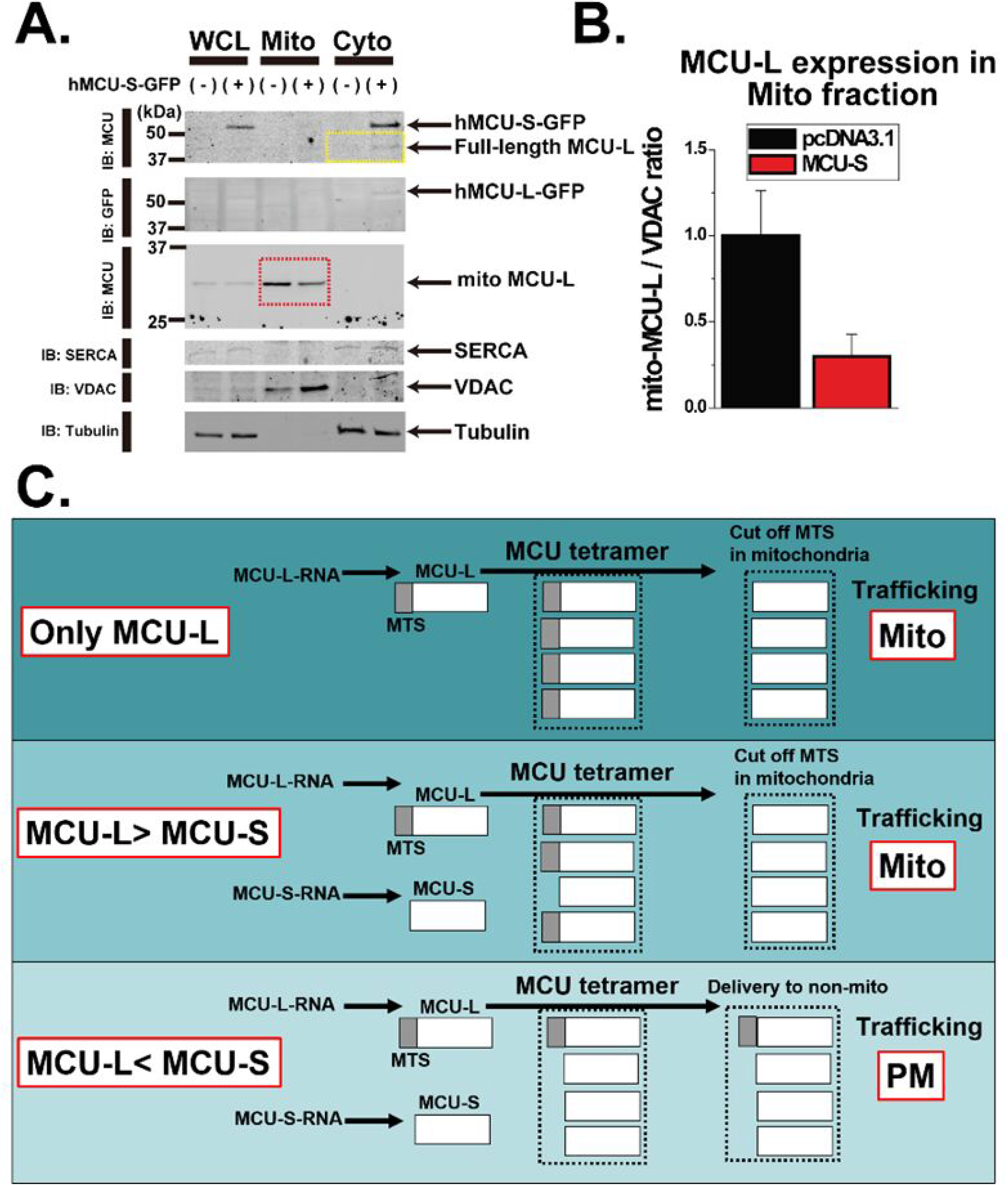
MCU-S inhibits mtCUC trafficking to mitochondria. **A.** Detecting human MCU variants in HEK293T cells overexpressing MCU-S-GFP or GFP (as a control) using fractionated protein. The mitochondria-enriched protein fraction (Mito) and cytosolic protein fraction (Cyto), which contains the ER proteins and PM proteins, were isolated as previously described with different centrifugation (see details in method section). Whole cell lysates (WCL) were also prepared for comparison. Tubulin was used as a loading control for WCL. SERCA and VDAC were used for loading and quality controls for Cyto and Mito fractions, respectively. The yellow-dotted box indicates the location of the band presumably corresponding to endogenous full-length MCU-L protein that includes the MTS (i.e., ∼40 kDa protein expressed in Cyto). The red-dotted box indicates the location of the bands presumably corresponding to endogenous MCU-L without MTS (i.e., ∼32 kDa protein expressed in the Mito). **B.** Summary data of A. **C.** Schematic diagram of the effect of MCU-S variant on MCU channel trafficking in the different MCU-L/MCU-S ratio. Oligomerization of ion-channel pore subunits mainly occurs in the ER and/or Golgi before delivery to the targeted organelle (Nagaya & Papazian, 1997) and the mtCUC pore is formed by MCU tetramer. If the MCU-L subunit number is larger than MCU-S within the MCU heteromeric tetramer (i.e., in the majority of cells/tissues under normal conditions, See Fig. 1), the majority of MCU proteins composing the tetramer possess MTS and the MCU tetramer will likely be trafficked to the mitochondria. If MCU-S expression is high, such as during MCU-S overexpression or in human platelets, MCU tetramers do not specifically traffic to mitochondria and may be sorted into other membrane structures, such as PM.

**Fig. EV3.**
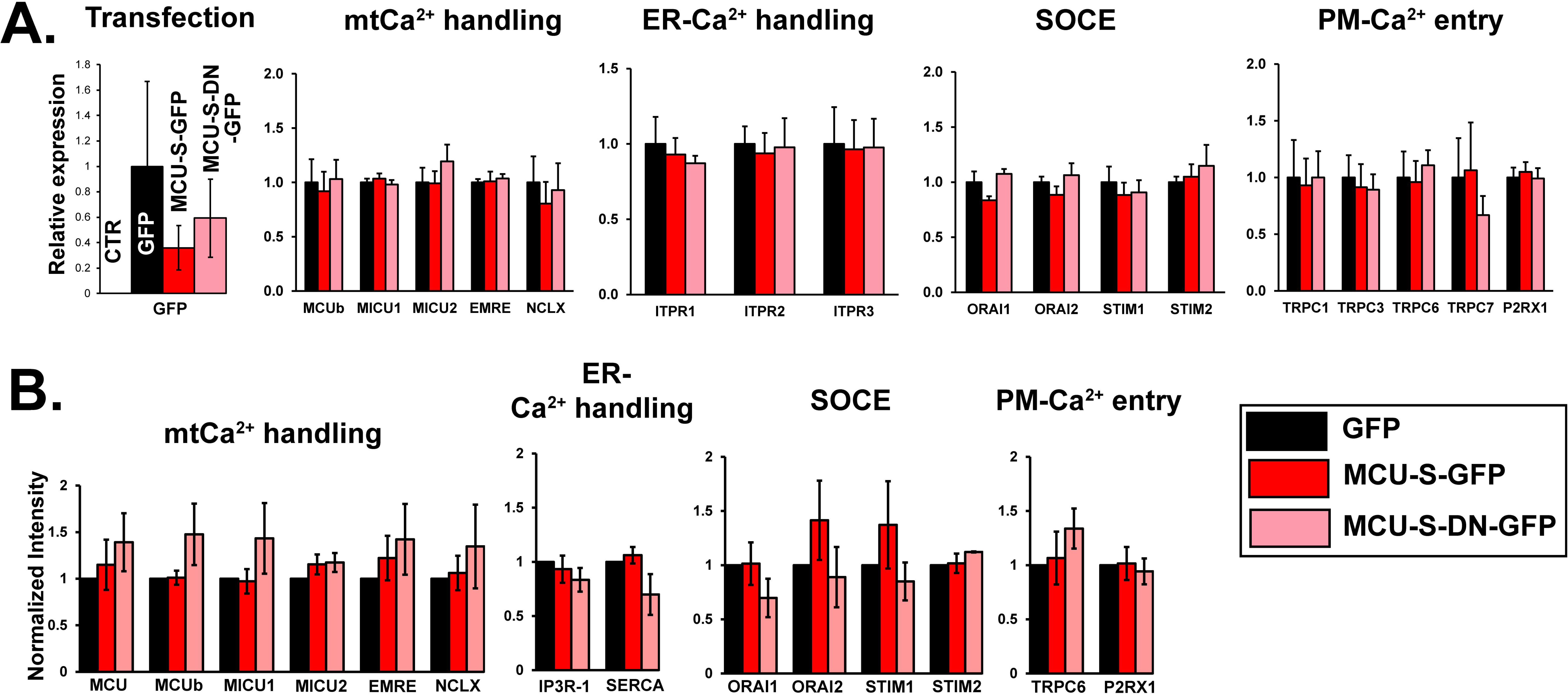
Effect of MCU-S expression on Ca^2+^ handling proteins in MEG-01 cells. **A.** Real-time qPCR analysis of major Ca^2+^ handling proteins in MEG-01 cells after transient transfection of MCU-S-GFP or MCU-S-DN-GFP. GFP was used as a control. CTR, non-transfected MEG-01 cells. N=3. **B.** Protein expression levels of major Ca^2+^ handling proteins in MEG-01 cells quantified by Western blotting. Tubulin was used as a loading control. All results were normalized to the value from the cells transfected with GFP. N=3 to 5.

**Fig. EV4.**
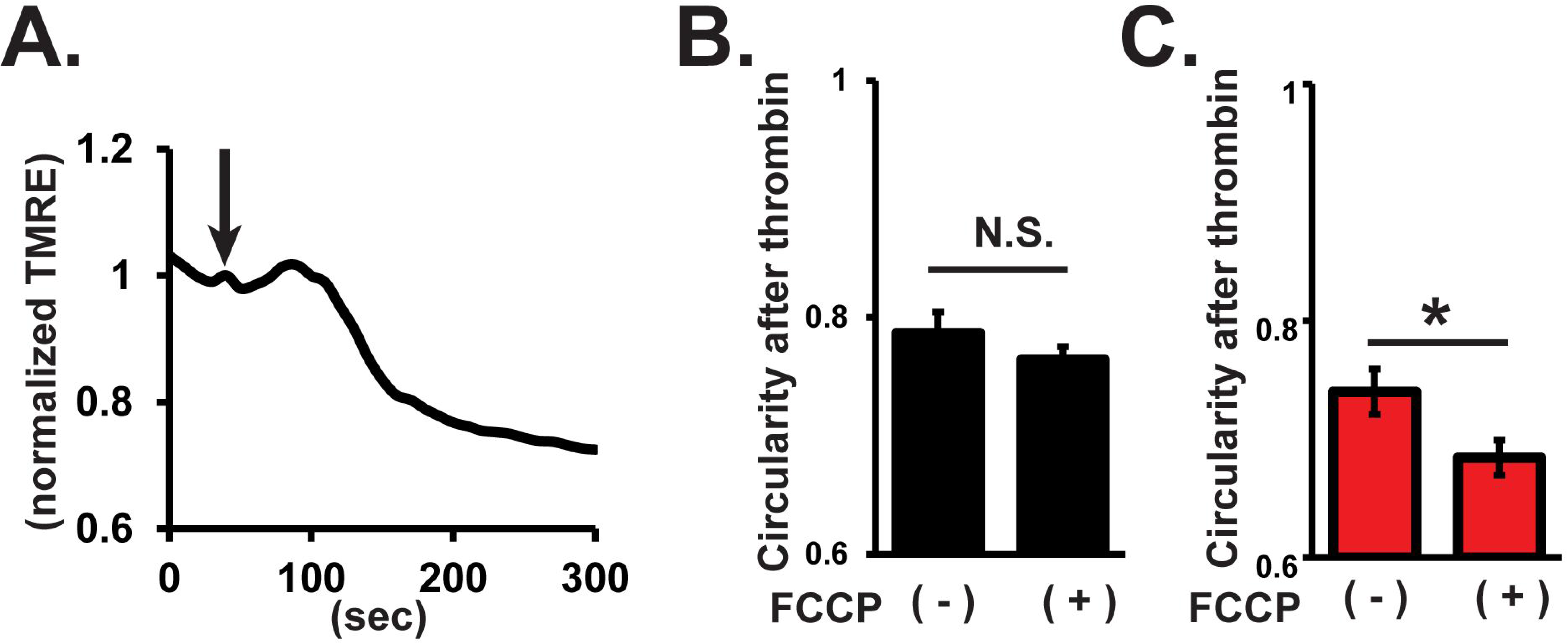
Effect of the mitochondrial uncoupler FCCP on thrombin-mediated morphological changes in MEG-01 cells. **A.** Representative trace of the changes in the fluorescence intensity of *Ψ_m_*-sensitive dye TMRE in response to treatment with 500 nM FCCP in a single MEG-01 cell. The fluorescence value at each time point was normalized with the value before FCCP treatment. **B.** Effect of FCCP on thrombin-mediated morphological changes in MEG-01 cells. MEG-01 cells transiently transfected with GFP were incubated with 500 nM for 5 minutes before thrombin stimulation. Morphological changes after thrombin stimulation were assessed glass-bottom micro dishes assessed by phase contrast images. Circularity reaches 1 if the cell is perfect circle and becomes low in complex cell shapes. N.S., not significant. **C.** Effect of FCCP on thrombin-mediated morphological changes in MEG-01 cells expressing MCU-S. MEG-01 cells transiently transfected with MCU-S-GFP were incubated with 500 nM FCCP for 5 minutes before thrombin stimulation. \**p*<0.05.

## References

Abbasian N, Millington-Burgess SL, Chabra S, Malcor J, & Harper MT (2020) Supramaximal calcium signaling triggers procoagulant platelet formation. Blood Adv 4: 154–164

Alevriadou BR, Patel A, Noble M, Ghosh S, Gohil VM, Stathopulos PB, & Madesh M (2021) Molecular nature and physiological role of the mitochondrial calcium uniporter channel. Am J Physiol Cell Physiol 320: C465–C482

Almagro Armenteros JJ, Salvatore M, Emanuelsson O, Winther O, von Heijne G, Elofsson A, & Nielsen H (2019) Detecting sequence signals in targeting peptides using deep learning. Life Sci Alliance 2: e201900429. doi: 10.26508/lsa.201900429. Print 2019 Oct

Antonipillai J, Mittelstaedt K, Rigby S, Bassler N, & Bernard O (2020) LIM kinase 2 (LIMK2) may play an essential role in platelet function. Exp Cell Res 388: 111822

Antonipillai J, Rigby S, Bassler N, Peter K, & Bernard O (2019) Pharmacological inhibition of LIM kinase pathway impairs platelets functionality and facilitates thrombolysis. Exp Cell Res 382: 111458

Baughman JM, Perocchi F, Girgis HS, Plovanich M, Belcher-Timme CA, Sancak Y, Bao XR, Strittmatter L, Goldberger O, Bogorad RL, Koteliansky V, & Mootha VK (2011) Integrative genomics identifies MCU as an essential component of the mitochondrial calcium uniporter. Nature 476: 341–345

Bearer EL, Prakash JM, & Li Z (2002) Actin dynamics in platelets. Int Rev Cytol 217: 137–182

Calvo SE, Clauser KR, & Mootha VK (2016) MitoCarta2.0: an updated inventory of mammalian mitochondrial proteins. Nucleic Acids Res 44: 1251

Calvo-Rodriguez M, Hou SS, Snyder AC, Kharitonova EK, Russ AN, Das S, Fan Z, Muzikansky A, Garcia-Alloza M, Serrano-Pozo A, Hudry E, & Bacskai BJ (2020) Increased mitochondrial calcium levels associated with neuronal death in a mouse model of Alzheimer’s disease. Nat Commun 11: 2146–2

Carninci P, Kasukawa T, Katayama S, Gough J, Frith MC, Maeda N, Oyama R, Ravasi T, Lenhard B, Wells C, Kodzius R, Shimokawa K, Bajic VB, Brenner SE, Batalov S, Forrest AR, Zavolan M, Davis MJ, Wilming LG, Aidinis V et al (2005) The transcriptional landscape of the mammalian genome. Science 309: 1559–1563

Chaudhuri D, Sancak Y, Mootha VK, & Clapham DE (2013) MCU encodes the pore conducting mitochondrial calcium currents. Elife 2: e00704

Claros MG & Vincens P (1996) Computational method to predict mitochondrially imported proteins and their targeting sequences. Eur J Biochem 241: 779–786

Dasgupta SK, Le A, Da Q, Cruz M, Rumbaut RE, & Thiagarajan P (2016) Wdr1-Dependent Actin Reorganization in Platelet Activation. PLoS One 11: e0162897

De Stefani D, Raffaello A, Teardo E, Szabo I, & Rizzuto R (2011) A forty-kilodalton protein of the inner membrane is the mitochondrial calcium uniporter. Nature 476: 336–340

Denorme F & Campbell RA (2022) Procoagulant platelets: novel players in thromboinflammation. Am J Physiol Cell Physiol 323: C951–C958

Dhenge A, Kuhikar R, Kale V, & Limaye L (2019) Regulation of differentiation of MEG01 to megakaryocytes and platelet-like particles by Valproic acid through Notch3 mediated actin polymerization. Platelets 30: 780–795

Dreos R, Ambrosini G, Perier RC, & Bucher P (2015) The Eukaryotic Promoter Database: expansion of EPDnew and new promoter analysis tools. Nucleic Acids Res 43: 92

Ghosh S, Basu Ball W, Madaris TR, Srikantan S, Madesh M, Mootha VK, & Gohil VM (2020) An essential role for cardiolipin in the stability and function of the mitochondrial calcium uniporter. Proc Natl Acad Sci U S A 117: 16383–16390

Hamilton S, Terentyeva R, Kim TY, Bronk P, Clements RT, O-Uchi J, Csordas G, Choi BR, & Terentyev D (2018) Pharmacological Modulation of Mitochondrial Ca(2+) Content Regulates Sarcoplasmic Reticulum Ca(2+) Release via Oxidation of the Ryanodine Receptor by Mitochondria-Derived Reactive Oxygen Species. Front Physiol 9: 1831

Harper MT & Poole AW (2011) Store-operated calcium entry and non-capacitative calcium entry have distinct roles in thrombin-induced calcium signalling in human platelets. Cell Calcium 50: 351–358

Heo Y, Jeon H, & Namkung W (2022) PAR4-Mediated PI3K/Akt and RhoA/ROCK Signaling Pathways Are Essential for Thrombin-Induced Morphological Changes in MEG-01 Cells. Int J Mol Sci 23: 776. doi: 10.3390/ijms23020776

Jhun BS, Mishra J, Monaco S, Fu D, Jiang W, Sheu SS, & O-Uchi J (2016) The mitochondrial Ca2+ uniporter: Regulation by auxiliary subunits and signal transduction pathways. Am J Physiol Cell Physiol 311(**1**): C67–C80

Jhun BS, O-Uchi J, Adaniya SM, Mancini TJ, Cao JL, King ME, Landi AK, Ma H, Shin M, Yang D, Xu X, Yoon Y, Choudhary G, Clements RT, Mende U, & Sheu SS (2018) Protein kinase D activation induces mitochondrial fragmentation and dysfunction in cardiomyocytes. J Physiol 596: 827–855

Jhun BS, O-Uchi J, Wang W, Ha CH, Zhao J, Kim JY, Wong C, Dirksen RT, Lopes CM, & Jin ZG (2012) Adrenergic signaling controls RGK-dependent trafficking of cardiac voltage-gated L-type Ca2+ channels through PKD1. Circ Res 110: 59–70

Kawai J, Shinagawa A, Shibata K, Yoshino M, Itoh M, Ishii Y, Arakawa T, Hara A, Fukunishi Y, Konno H, Adachi J, Fukuda S, Aizawa K, Izawa M, Nishi K, Kiyosawa H, Kondo S, Yamanaka I, Saito T, Okazaki Y et al (2001) Functional annotation of a full-length mouse cDNA collection. Nature 409: 685–690

Kholmukhamedov A, Janecke R, Choo H, & Jobe SM (2018) The mitochondrial calcium uniporter regulates procoagulant platelet formation. J Thromb Haemost 16: 2315–2321

Kirichok Y, Krapivinsky G, & Clapham DE (2004) The mitochondrial calcium uniporter is a highly selective ion channel. Nature 427: 360–364

Lee CW, Vitriol EA, Shim S, Wise AL, Velayutham RP, & Zheng JQ (2013) Dynamic localization of G-actin during membrane protrusion in neuronal motility. Curr Biol 23: 1046–1056

Mammadova-Bach E, Nagy M, Heemskerk JWM, Nieswandt B, & Braun A (2019) Store-operated calcium entry in thrombosis and thrombo-inflammation. Cell Calcium 77: 39–48

Mammucari C, Raffaello A, Vecellio Reane D, Gherardi G, De Mario A, & Rizzuto R (2018) Mitochondrial calcium uptake in organ physiology: from molecular mechanism to animal models. Pflugers Arch

Millington-Burgess SL & Harper MT (2021) Cytosolic and mitochondrial Ca(2+) signaling in procoagulant platelets. Platelets 32: 855–862

Miura K, Fujibuchi W, & Unno M (2012) Splice variants in apoptotic pathway. Exp Oncol 34: 212–217

Nagaya N & Papazian DM (1997) Potassium channel alpha and beta subunits assemble in the endoplasmic reticulum. J Biol Chem 272: 3022–3027

Okazaki Y, Furuno M, Kasukawa T, Adachi J, Bono H, Kondo S, Nikaido I, Osato N, Saito R, Suzuki H, Yamanaka I, Kiyosawa H, Yagi K, Tomaru Y, Hasegawa Y, Nogami A, Schonbach C, Gojobori T, Baldarelli R, Hill DP et al (2002) Analysis of the mouse transcriptome based on functional annotation of 60,770 full-length cDNAs. Nature 420: 563–573

O-Uchi J, Jhun BS, Hurst S, Bisetto S, Gross P, Chen M, Kettlewell S, Park J, Oyamada H, Smith GL, Murayama T, & Sheu SS (2013) Overexpression of ryanodine receptor type 1 enhances mitochondrial fragmentation and Ca2+-induced ATP production in cardiac H9c2 myoblasts. Am J Physiol Heart Circ Physiol

O-Uchi J, Jhun BS, Xu S, Hurst S, Raffaello A, Liu X, Yi B, Zhang H, Gross P, Mishra J, Ainbinder A, Kettlewell S, Smith GL, Dirksen RT, Wang W, Rizzuto R, & Sheu SS (2014) Adrenergic signaling regulates mitochondrial Ca2+ uptake through Pyk2-dependent tyrosine phosphorylation of the mitochondrial Ca2+ uniporter. Antioxid Redox Signal 21: 863–879

Patron M, Checchetto V, Raffaello A, Teardo E, Vecellio Reane D, Mantoan M, Granatiero V, Szabo I, De Stefani D, & Rizzuto R (2014) MICU1 and MICU2 finely tune the mitochondrial Ca2+ uniporter by exerting opposite effects on MCU activity. Mol Cell 53: 726–737

Raffaello A, De Stefani D, Sabbadin D, Teardo E, Merli G, Picard A, Checchetto V, Moro S, Szabo I, & Rizzuto R (2013) The mitochondrial calcium uniporter is a multimer that can include a dominant-negative pore-forming subunit. EMBO J 32: 2362–2376

Schindelin J, Arganda-Carreras I, Frise E, Kaynig V, Longair M, Pietzsch T, Preibisch S, Rueden C, Saalfeld S, Schmid B, Tinevez JY, White DJ, Hartenstein V, Eliceiri K, Tomancak P, & Cardona A (2012) Fiji: an open-source platform for biological-image analysis. Nat Methods 9: 676–682

Strausberg RL, Feingold EA, Grouse LH, Derge JG, Klausner RD, Collins FS, Wagner L, Shenmen CM, Schuler GD, Altschul SF, Zeeberg B, Buetow KH, Schaefer CF, Bhat NK, Hopkins RF, Jordan H, Moore T, Max SI, Wang J, Hsieh F et al (2002) Generation and initial analysis of more than 15,000 full-length human and mouse cDNA sequences. Proc Natl Acad Sci U S A 99: 16899–16903

Untergasser A, Cutcutache I, Koressaar T, Ye J, Faircloth BC, Remm M, & Rozen SG (2012) Primer3--new capabilities and interfaces. Nucleic Acids Res 40: e115

Vang A, da Silva Goncalves Bos D, Fernandez-Nicolas A, Zhang P, Morrison AR, Mancini TJ, Clements RT, Polina I, Cypress MW, Jhun BS, Hawrot E, Mende U, O-Uchi J, & Choudhary G (2021) alpha7 Nicotinic acetylcholine receptor mediates right ventricular fibrosis and diastolic dysfunction in pulmonary hypertension. JCI Insight 6: 10.1172/jci.insight.142945

Varga-Szabo D, Braun A, & Nieswandt B (2009) Calcium signaling in platelets. J Thromb Haemost 7: 1057–1066

Xie A, Song Z, Liu H, Zhou A, Shi G, Wang Q, Gu L, Liu M, Xie LH, Qu Z, & Dudley SC,Jr (2018) Mitochondrial Ca(2+) Influx Contributes to Arrhythmic Risk in Nonischemic Cardiomyopathy. J Am Heart Assoc 7: 10.1161/JAHA.117.007805

Yoon Y, Krueger EW, Oswald BJ, & McNiven MA (2003) The mitochondrial protein hFis1 regulates mitochondrial fission in mammalian cells through an interaction with the dynamin-like protein DLP1. Mol Cell Biol 23: 5409–5420

Zaglia T, Ceriotti P, Campo A, Borile G, Armani A, Carullo P, Prando V, Coppini R, Vida V, Stolen TO, Ulrik W, Cerbai E, Stellin G, Faggian G, De Stefani D, Sandri M, Rizzuto R, Di Lisa F, Pozzan T, Catalucci D et al (2017) Content of mitochondrial calcium uniporter (MCU) in cardiomyocytes is regulated by microRNA-1 in physiologic and pathologic hypertrophy. Proc Natl Acad Sci U S A 114: E9006–E9015

